# Explorations of using a convolutional neural network to understand brain activations during movie watching

**DOI:** 10.1101/2024.01.20.576341

**Authors:** Wonbum Sohn, Xin Di, Zhen Liang, Zhiguo Zhang, Bharat B. Biswal

## Abstract

Neuroimaging studies increasingly use naturalistic stimuli like video clips to trigger complex brain activations, but the complexity of such stimuli makes it difficult to assign specific functions to the resulting brain activations, particularly for higher-level content like social interactions. To address this challenge, researchers have turned to deep neural networks, e.g., convolutional neural networks (CNNs). CNNs have shown success in image recognition due to their different levels of features enabling high performance. In this study, we used pre-trained VGG-16, a popular CNN model, to analyze video data and extract hierarchical features from low-level shallow layers to high-level deeper layers, linking these activations to different levels of activation of the human brain. We hypothesized that activations in different layers of VGG-16 would be associated with different levels of brain activation and visual processing hierarchy in the brain. We were also curious about which brain regions would be associated with deeper convolutional layers in VGG-16. The study analyzed a functional MRI (fMRI) dataset where participants watched the cartoon movie Partly Cloudy. Frames of the videos were fed into VGG-16, and activation maps from different kernels and layers were extracted. Time series of the average activation patterns for each kernel were created and fed into a voxel-wise model to study brain activations. Results showed that lower convolutional layers (1^st^ convolutional layer) were mostly associated with lower visual regions, but some kernels (6, 19, 24, 42, 55, and 58) surprisingly showed associations with activations in the posterior cingulate cortex, part of the default mode network. Deeper convolutional layers were associated with more anterior and lateral portions of the visual cortex (e.g., the lateral occipital complex) and the supramarginal gyrus. Analyzing activation features associated with different brain regions showed the promise and limitations of using CNNs to link video content to brain functions.

## 1. Introduction

Recently, naturalistic stimuli, like movies and stories, have increasingly been employed to study brain functions in human neuroimaging research. This approach offers several advantages over traditional task-based fMRI experiments. One of the primary benefits is that naturalistic stimuli closely resemble real-life situations, enabling the elicitation of complex cognitive processes. On the other hand, compared with resting-state fMRI, naturalistic stimuli allow for a higher level of experimental control, resulting in improved participant cooperation and increased reliability of research findings. A pivotal study by Hasson et al. demonstrated that different participants watching the same video stimulus exhibited similar patterns of brain activity across distributed brain regions (Hasson et al., 2004). This finding led to the widespread use of inter-subject correlation as a means to identify activity and connectivity patterns associated with various stimuli (Chen et al., 2020; Di & Biswal, 2020; Nastase, 2019). Despite these advancements, one major challenge is linking the observed brain data to the contents of naturalistic stimuli, such as videos and audios, due to their inherent complexity.

Numerous analytical approaches have emerged to study the complexities of naturalistic stimuli. One conventional method is utilizing human subjective ratings. For instance, researchers have asked participants to rate their perceived motion while watching a cartoon video and then employed general linear model to map brain responses related to motion perception. This approach identified motion-sensitive brain regions in the middle temporal lobe (Rao et al., 2007). Furthermore, subjective affective states can be reported and linked to brain activations and dynamic connectivity (Raz et al., 2012; Sun et al., 2022). Another approach is manually tagging objects of interest to investigate category-specific brain activations (Richardson et al., 2018). Advancements in machine learning technologies have also been leveraged. Previous studies used traditional computer vision models to extract global motion, local motion, and residual models based on motion flow and patterns from videos. They found that the medial posterior parietal cortex, V5+, and V1∼V4 were activated in the scenes of the global motion model, local motion model, and residual model, respectively (Bartels et al., 2008). Celik et al. have built encoding models of various objects (car, bridge, etc.) from video stimuli to study category representations in the cerebral cortex (Çelik et al., 2021). Recently, convolutional neural networks (CNNs) have been used to extract visual features of videos, particularly in the context of face processing (Hu et al., 2023; Jiahui et al., 2022).

The naturalistic stimuli have been selected to explore intricate social functions like the theory of mind and empathy (Richardson et al., 2018). However, the field still lacks machine learning models that can effectively describe various aspects of social functions due to the complexity of the naturalistic stimuli. A recent study by McMahon and colleagues employed multiple machine learning models to extract different levels of features from videos containing social interactions (McMahon et al., 2023). They established a hierarchy of social interactions, primarily linked to the temporal lobe regions. Nevertheless, the higher-level features in their hierarchy still rely on manually selected features. In our current study, we aim to test the hypothesis of whether we can extract features related to social functions using convolutional neural networks. We systematically investigate how different convolutional layers are associated with the hierarchy of various brain regions. This approach may offer valuable insights into understanding the neural basis of social interactions and potentially uncover novel findings that were previously limited by manual feature selection methods.

Convolutional neural networks (CNNs) have demonstrated remarkable success in computer vision (Krizhevsky et al., 2017; Simonyan & Zisserman, 2015). One of fundamental elements in CNNs is the convolutional kernels, which extract local features from data. As the data progresses through deeper layers of convolutional kernels, more complex features are extracted. While CNNs are typically trained on large-scale image datasets for image recognition, encompassing 1,000 categories (Deng et al., 2009), we posit that a CNN trained on the ImageNet dataset might learned information relevant to social interactions. To explore this hypothesis, we investigate how features extracted from various kernels of convolutional layers correlate with brain activations in different brain regions. In a recent study, Hu and colleagues utilized the a pre-trained VGG-16 CNN to extract features from different layers while analyzing affective videos (Hu et al., 2023). They discovered that brain microstates calculated from electroencephalogram data were only correlated with features from deeper convolutional layers (layers 11, 12, and 13). Building upon this work, our current study employs fMRI data, which provides superior spatial resolution. This enables us to examine how different brain regions are associated with the features extracted from diverse kernels of convolutional layers. By leveraging the strengths of fMRI, we aim to gain deeper insights into the relationship between neural activations and the hierarchical visual representations generated by CNNs.

In this study, we employed a single pre-trained VGG-16 network, one of image feature extractors, to analyze a short, animated movie and extract features at different levels from the convolutional layers. Our primary aim was to investigate how and where these diverse levels of features are represented in the human brain. To accomplish this, we collected fMRI data from young adult volunteers while they watched the same movie clip. We utilized a generalized linear model approach to map brain regions whose temporal activity pattern matched the feature activity pattern from specific kernels of a convolutional layer. Our hypothesis revolves around the notion that brain activation patterns will exhibit a hierarchy from low-level visual areas to high-level areas related to social interaction and empathy through CNN’s hierarchical feature maps. Of particular interest to us were the brain regions associated with higher convolutional layers. Considering that VGG-16 was trained for image classification, we postulated that higher convolutional layers might primarily represent categorical features, possibly linking to the ventral visual pathway in the brain. In addition, due to the rich image context in the movie, it could also encompass action, motion, and social information, which may be more prevalent in deeper convolutional layers of VGG-16. We were especially intrigued by whether the dorsal visual pathway, and even the third "social" pathway, could also be associated with features from deeper layers. If this proves to be true, VGG-16 could serve as a valuable tool for studying social functions using video stimuli, extending beyond its conventional image classification capabilities.

## 2. Materials and methods

### 2.1. fMRI Data

The fMRI data were obtained from openneuro (https://openneuro.org/; accession #: ds000228). Only adults’ data (n=33) were used for this study. The effective sample consisted of 17 females and 12 males (age range = 18-39 years, mean = 24.6, SD = 5.3), excluding subjects with excessive head motion or poor coverage according to the criteria mentioned in the previous study (Di & Biswal, 2020).

During the fMRI acquisition, all subjects watched "Partly Cloudy" animation of the Pixar (https://www.pixar.com/partly-cloudy#par-tly-cloudy-1) for about 6 minutes with a black screen of first 10 seconds (1 - 5 TRs) (Richardson et al., 2018). Structural and functional MRI data were acquired by a 3 Tesla Siemens Tim Trio scanner at the Massachusetts Institute of Technology. The T1-weighted structural MRI data were gathered in 176 interleaved sagittal slices with 1mm isotropic. All fMRI data were acquired by a gradient-echo EPI (repetition time (TR) = 2s, echo time (TE) = 30ms, flip angle = 90 deg). 168 fMRI images were measured from each subject.

The fMRI data were analyzed using SPM12 and MATLAB (R2021b) as described in our previous study (Di & Biswal, 2020). Briefly, first, the anatomical T1 images of all subjects were segmented into six segments. Afterward, all skulls were removed from the T1 images. All functional images were realigned concerning the first image. In this process, the degree of translation and rotation was calculated, and subjects with a maximum framewise displacement greater than 1.5 mm or 1.5 deg were removed (Di & Biswal, 2015). The remaining functional images were coregistered with the skull-stripped anatomical image of the same subject. Next, the anatomical and functional images were normalized to MNI space, and the voxel size was resampled to 3 x 3 x 3 *mm*^3^. Finally, the functional images were spatially smoothed through an 8mm Gaussian kernel.

### 2.2. Video analysis

Given the TR of 2 seconds, each fMRI image corresponds to 2 seconds of video contents. Therefore, we averaged the video frames for every 2 seconds before extracting features from the video. Since the frame rate was 24 frames/s, 48 frames were averaged into one image. As a result, the 8370 total frames of the video clip were converted to 175 images.

We used the pre-trained VGG-16 model to process the image data. VGG-16 consists of 13 convolutional layers, 5 max-pooling layers, and 3 fully-connected layers (Figure 1A). Each averaged image with 720 x 1280 x 3 dimension was fed to the VGG-16 model. The input images were convolved with 64 3 x 3 x 3 kernels in the 1^st^ convolutional layer and then generated 720 x 1280 x 64 feature maps. Feature maps show the location and intensity of various patterns contained in the input. These were used as an input of the 2^nd^ convolutional layer and convolved with 64 3 x 3 x 64 kernels, generating 720 x 1280 x 64 feature maps. Afterward, the size of the feature maps was reduced to 360 x 640 x 64 through the max-pooling layer. This process has repeated 5 times, and finally, the size of the feature maps became 45 x 80 x 512. The last max-pooling layer and the three fully-connected layers of VGG-16 were not used because this step aimed to extract feature maps from the averaged input images instead of classification. Each layer receives the output feature maps of the previous layer as input and generates higher-level feature maps. As a result, low-level features such as edges, lines, and similar color regions were detected in shallow convolutional layers. On the other hand, abstract and synthetic high-level features were shown in the deeper layers.

**Figure 1.**
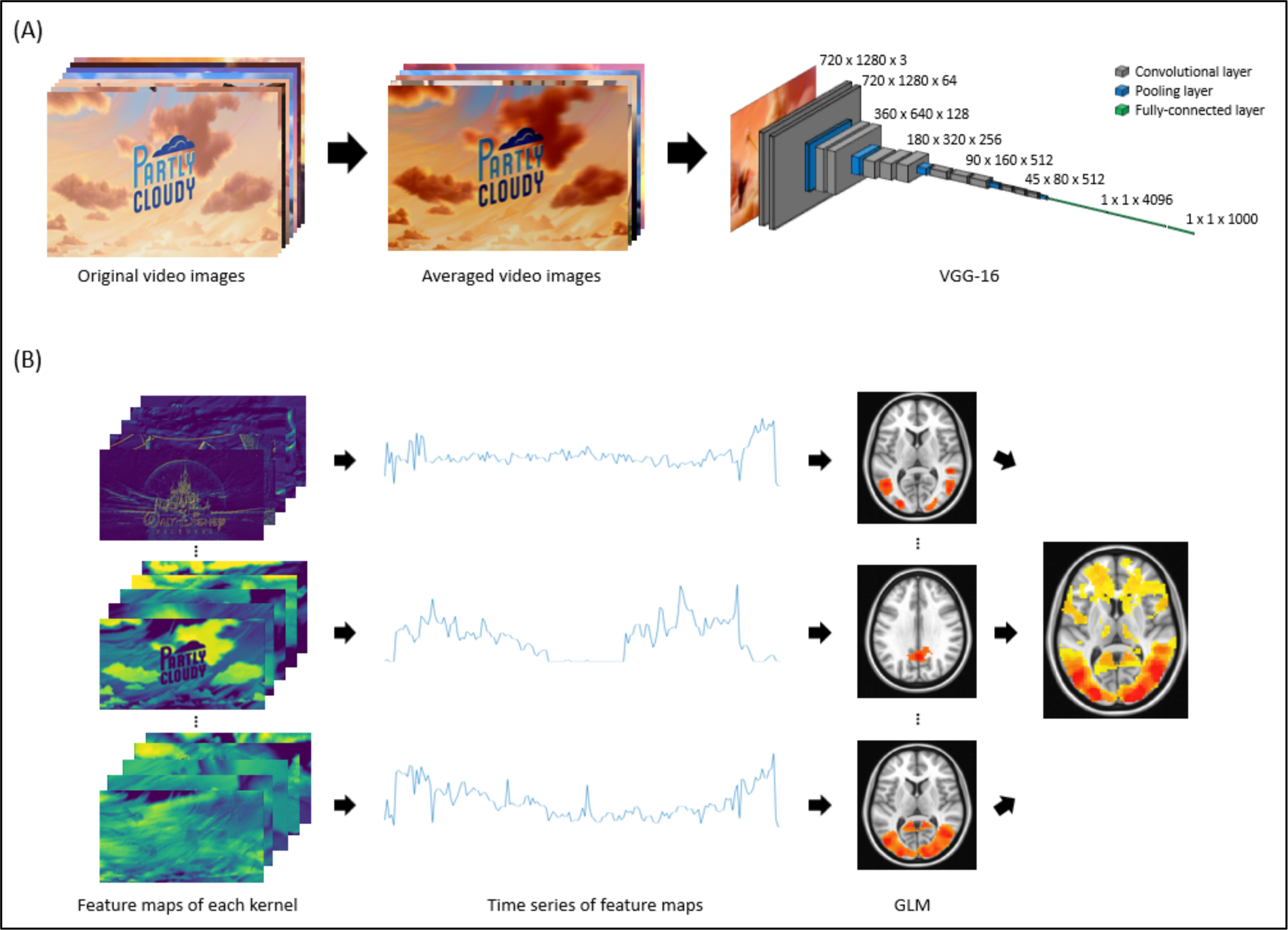
Illustration of feature extraction from the video and GLM. (A) 8,370 images of the original video were converted into 175 images based on TR 2s and the 175 images were used in VGG-16 for feature extraction. (B) 2D feature maps from each kernel were transformed into time series, and then the time series was used for GLM on the fMRI data. Kernel activation maps were calculated by averaging the activation maps of every kernel of a convolutional layer in VGG-16.

### 2.3. Linking VGG-16 activations to brain activations

For each TR, each kernel of a convolutional layer of the VGG-16 model generates a distinct activation map. In the context of general linear model (GLM) analysis—which scrutinizes brain regions exhibiting patterns akin to a specific time series during the measurement period—each activation map was averaged into a single value. Consequently, with 175 input images sampling the entire video, a 1D time series of 175 time was generated for each kernel. This resulted in 64 time series for 1^st^ and 2^nd^ convolutional layers, and 512 time series for 11^th^, 12^th^, and 13^th^ convolutional layers. The size of the 1D time series for each convolutional layer was detailed in Table 1. Here we used the averaged activations to match gross neuronal activities that are measured by fMRI (Figure 1B).

In order to select two convolutional layers with a large difference between feature maps, we calculated the correlation between layers. We averaged the 1D time series data that exist as many as the number of kernels in each convolutional layer to obtain one averaged 1D time series data for each layer. We selected the 1^st^ and 13^th^ convolutional layers with the lowest correlation (r = -0.054) for GLM and reverse analysis.

We performed voxel-wise GLM using the 1D time series of the kernels and preprocessed fMRI data. First, the first 5 TRs of the fMRI data obtained from the black screen were excluded because they were unrelated to the video. Next, to match with fMRI data length, the 1D time series were used only for the first 163 TRs. Friston’s 24-parameter model was also added as a regressor to consider the effect of head motion (Friston et al., 1996). To check the brain regions activated by the 1D time series of video features, the contrast was set to 1 for only the video time series and 0 for the remaining Friston’s 24-parameter model parts. Finally, a group-level one-sample t-test was performed for each kernel to confirm the brain networks commonly activated by the features across the subjects. After that, the cluster of the GLM results was performed based on p<0.001. After that, the group-level one-sample t-test results of all kernels for each convolutional layer were added to see which functional network of the brain was related as the convolutional layer deepened.

### 2.4. Analyze feature maps

In the voxel-wise GLM analysis, we identified brain regions that were activated by features extracted from different layers of VGG-16. We then performed reverse analysis to find patterns of feature maps related to the brain activations. Specially, we were interested in the brain activations in the default mode network and visual cortex prominent from features of the 1^st^ convolutional layer and in the supramarginal gyrus and lateral occipital complex regions prominent from features of the 13^th^ convolutional layer.

In the 1^st^ convolutional layer, the brain’s functional network obtained from the GLM with the kernel 6 related to the default mode network, especially posterior cingulate cortex, was masked with 0 and 1 to create a posterior cingulate mask. The generated posterior cingulate mask was applied to the preprocessed fMRI data of 29 subjects, leaving only the intensities of the brain regions related to the posterior cingulate cortex and converting the intensities of the other regions to 0. The intensities of the remaining posterior cingulate regions were averaged with one value for each TR time and then made time series with a length of 165 TRs for each subject. To determine the TR time at which the posterior cingulate region is actively activated, the time series ensemble of 29 subjects was averaged into one averaged fMRI time series. We identified the TR times with peaks in the averaged fMRI time series. Considering that the delay until the BOLD signal was generated after stimulation was 2 TR (4 seconds), feature maps of the kernel 6 corresponding to the TR time after subtracting 2 TRs from the identified TR time were extracted. We repeated this process for other kernels 19, 24, 42, 55, and 58 related to the posterior cingulate to extract related feature maps. We investigated whether common patterns exist in the extracted feature maps.

The same method was applied to kernels 7, 43, and 63 of the 1^st^ convolutional layer related to the visual cortex. In the 13^th^ convolutional layer, kernels 9, 43, and 128 related to lateral occipital complex and kernels 42, 103, 168, 276, 428, and 510 associated with supramarginal gyrus were confirmed for the reverse analysis. After extracting the relevant feature maps, we analyzed whether there were common patterns in the feature maps for each brain region.

## 3. Results

### 3.1. Variability of video time series with layer depth

We examined the variability between the 1D time series data obtained from each kernel in the convolutional layers and the correlation between the time series data across the convolutional layers. Principal component analysis (PCA) was used to maximize the variability of kernels in each convolutional layer. In the shallow layers (1^st^ ∼ 7^th^ convolutional layers), there was a huge difference in the proportion of PC1 representing the features variability depending on the layer (Figure 2A). On the other hand, in the deep layers (8^th^ ∼ 12^th^ convolutional layers), there was little difference in the variability proportion of PC1 according to the layer except for the 13^th^ convolutional layer. Overall, as the layer deepened, the proportion of PC1∼2 in variability tended to decrease. This means that more various features were extracted in deeper convolutional layers. To see the correlation between layers, a total of 13 averaged 1D time series data were used to calculate correlation coefficients (Figure 2B). Correlation coefficients between adjacent convolutional layers were high. On the other hand, the correlation coefficients were lower as the distance of the convolutional layers increased. This was because CNN receives the output of the previous convolutional layer as an input and performs a convolution operation.

**Figure 2.**
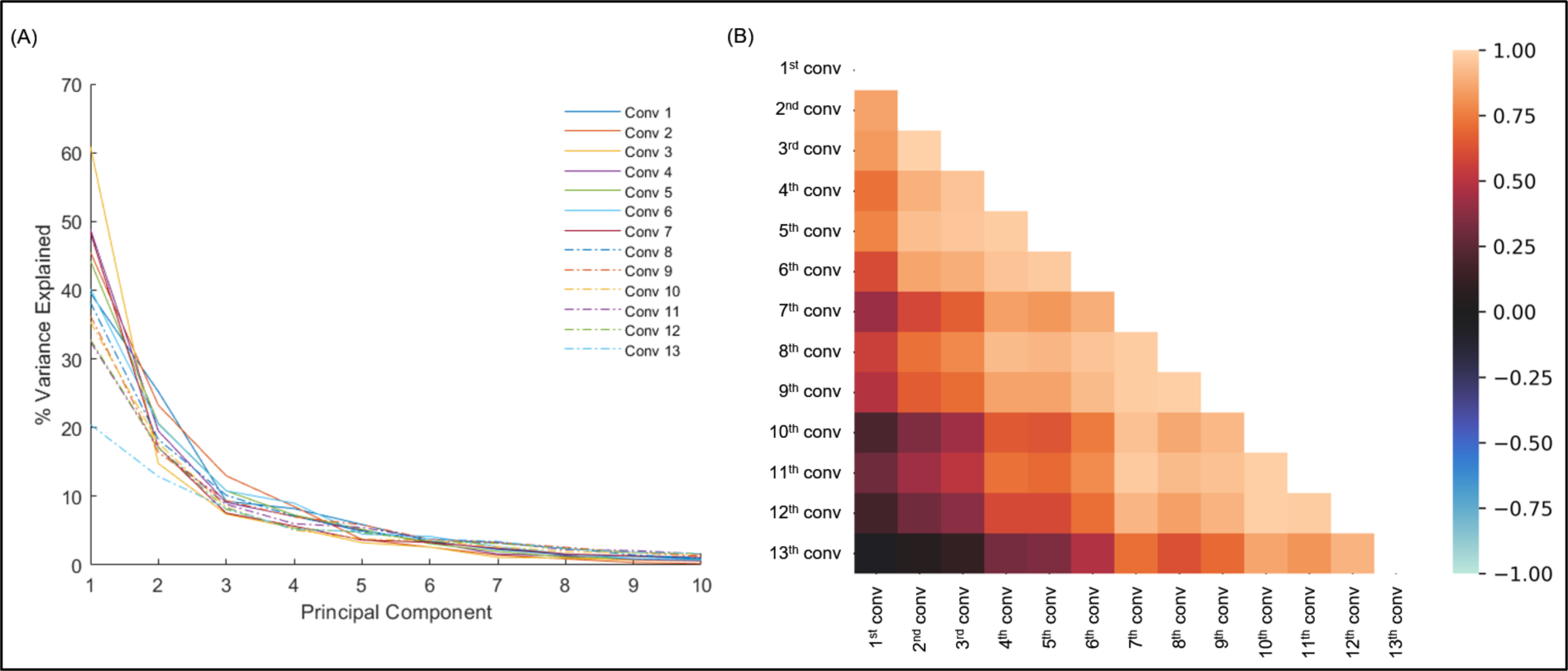
The characteristics of the convolutional layers. (A) The variance explained by the first ten principal components for the 13 convolutional layers. The variance percentage of PC1∼2 became lower when the convolutional layers went deeper. (B) A correlation matrix of averaged time series across the 13 convolutional layers. The correlation coefficient was higher between adjacent layers and lower between distant layers.

### 3.2. GLM results

After obtaining time series for each kernel in different convolutional layers, we performed voxel-wise analysis with brain activations measured with fMRI. Most of the GLMs produced statistically significant activations in various brain regions. We examined the distributions of activations in the brain in different convolutional layers of VGG-16 (Figure 3). There were widespread activations associated with different kernels in all the layers. The visual cortex was more likely to be activated in all the convolutional layers. But the activation patterns outside the visual cortex conveyed a shift in different convolutional layers. Specifically, the posterior cingulate cortex was more likely to be activated in shallow layers such as the 1^st^ and 3^rd^ convolutional layers. As layers went deeper, bilateral temporal and parietal regions were more likely to be activated.

**Figure 3.**
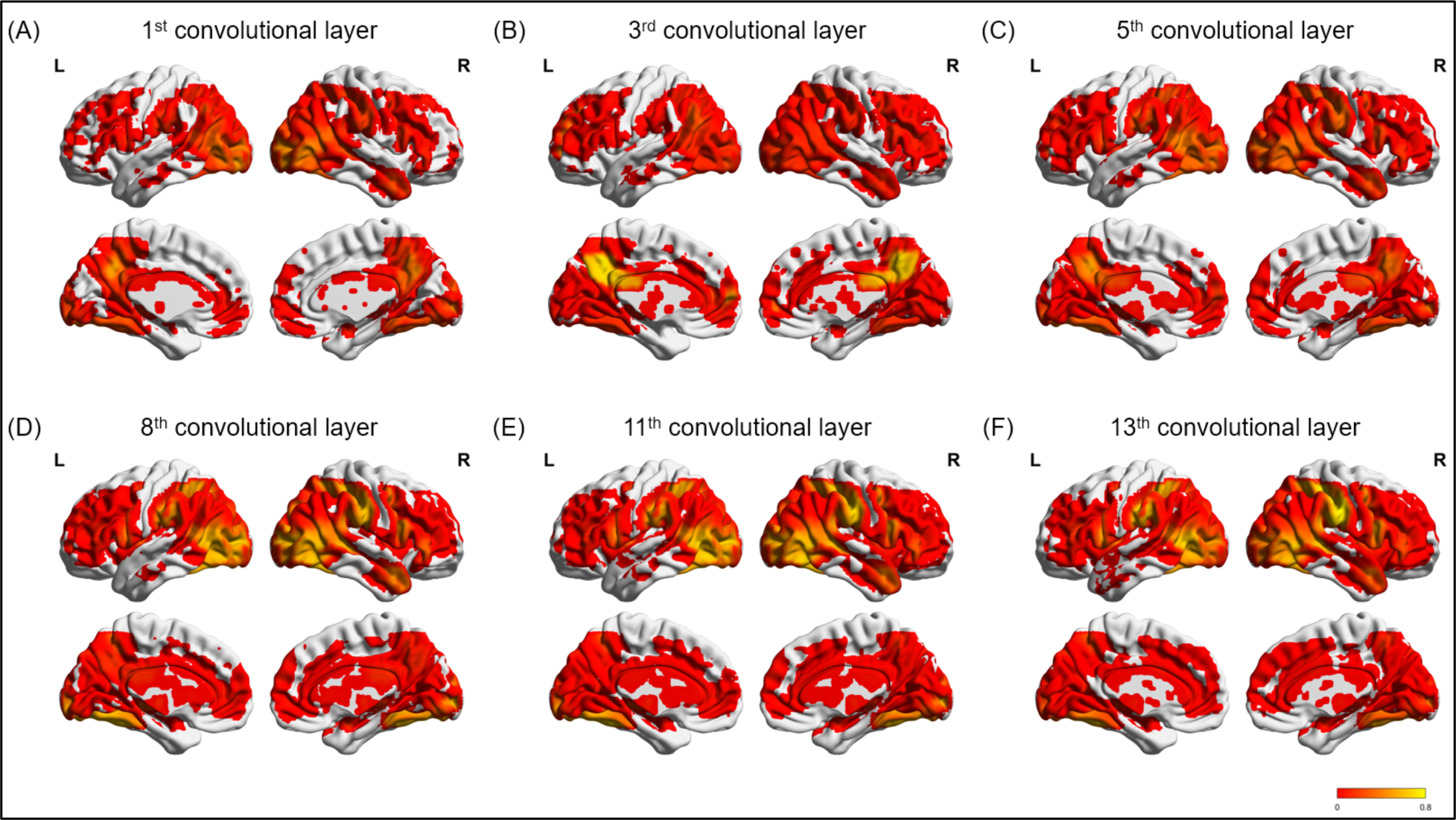
Activation probability by feature activations of the video in different kernels in the convolutional layers of 1^st^, 3^rd^, 5^th^, 8^th^, 11^th^, and 13^th^ in VGG-16. Binary activation maps for each kernel were averaged within a layer to form the probability map, resulting in a range between 0 to 1.

We contrasted the brain activation probability maps between the shallowest layer (1^st^ convolutional layer) and the deepest layer (13^th^ convolutional layer). Figure 4 clearly shows the different activation distributions between the two layers. The 1^st^ convolutional layer was more likely to be associated with the posterior visual cortex, as well as the posterior cingulate cortex, bilateral angular gyrus, and medial prefrontal cortex, which formed the default mode network. In contrast, the 13^th^ convolutional layer was more likely to be associated with supramarginal gyrus, lateral occipital complex, and superior temporal sulcus.

**Figure 4.**
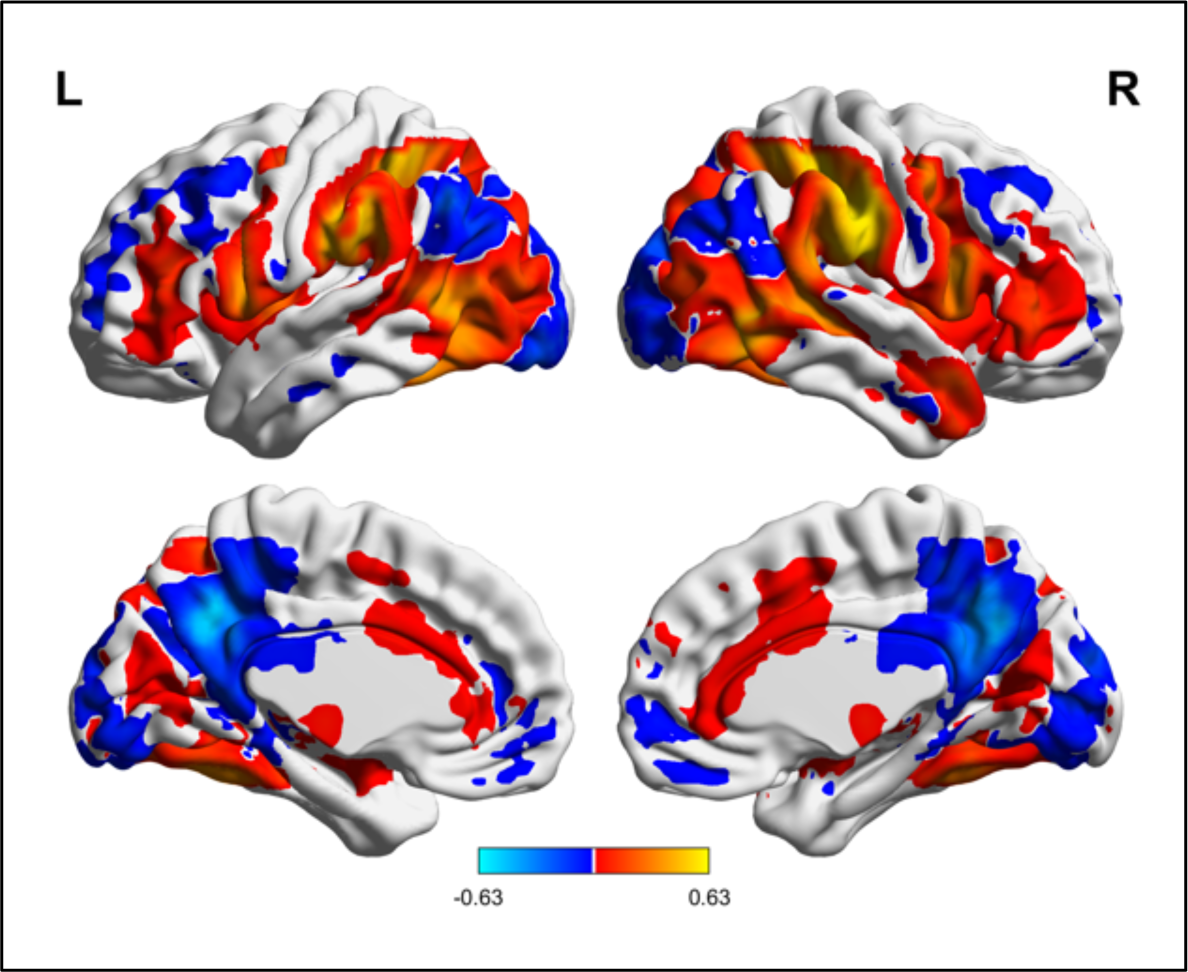
Differences in activation probability by feature activations in different kernels between the 13^th^ and the 1^st^ layers. Red is more active brain regions in the 13^th^ convolutional layer, and blue is more active brain regions in the 1^st^ convolutional layer.

### 3.3. Patterns in feature maps

To gain insights into the relationship between video features and brain activations across different regions, we conducted a reverse analysis. As a specific case, we focused on the brain regions that exhibited a higher propensity for activation based on features derived from the 1^st^ and 13^th^ convolutional layers. Specifically, we examined the visual cortex and posterior cingulate cortex region, which demonstrated a greater likelihood of activation in response to features from the 1^st^ convolutional layer (Figure 5). Additionally, we investigated the supramarginal gyrus and lateral occipital complex region, which exhibited a heightened probability of activation in response to features from the 13th convolutional layer (Figure 6). To investigate independent relationships between specific brain regions and features, kernels that simultaneously activate multiple brain regions were excluded.

**Figure 5.**
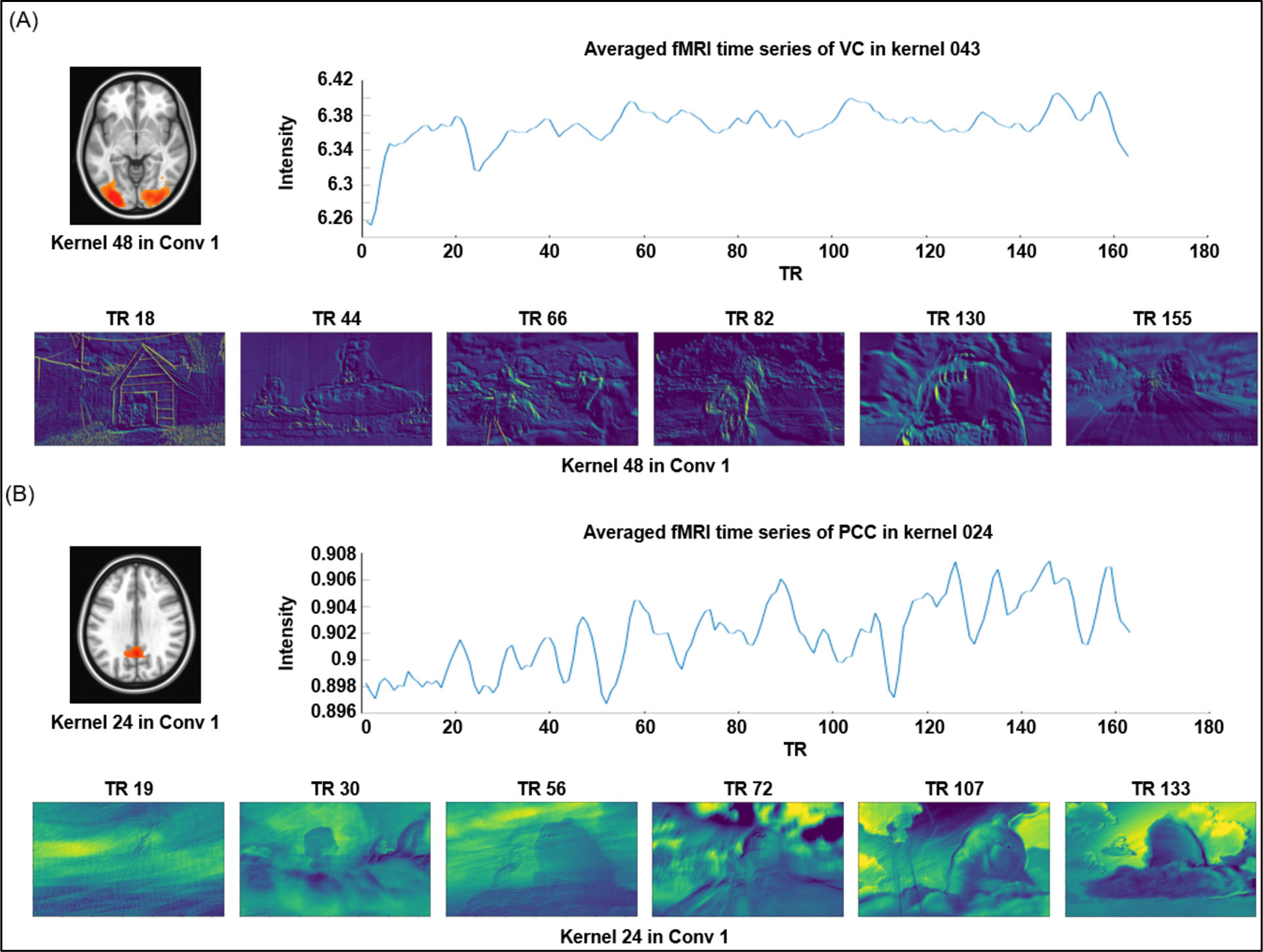
Reverse analysis of (A) the visual cortex and (B) posterior cingulate cortex. Activation maps were obtained as GLM results for (A) kernel 43 related to visual cortex and (B) kernel 24 related to posterior cingulate cortex. The averaged fMRI time series are for each brain region, and the feature maps are for the TR times associated with the peaks of the averaged fMRI time series.

**Figure 6.**
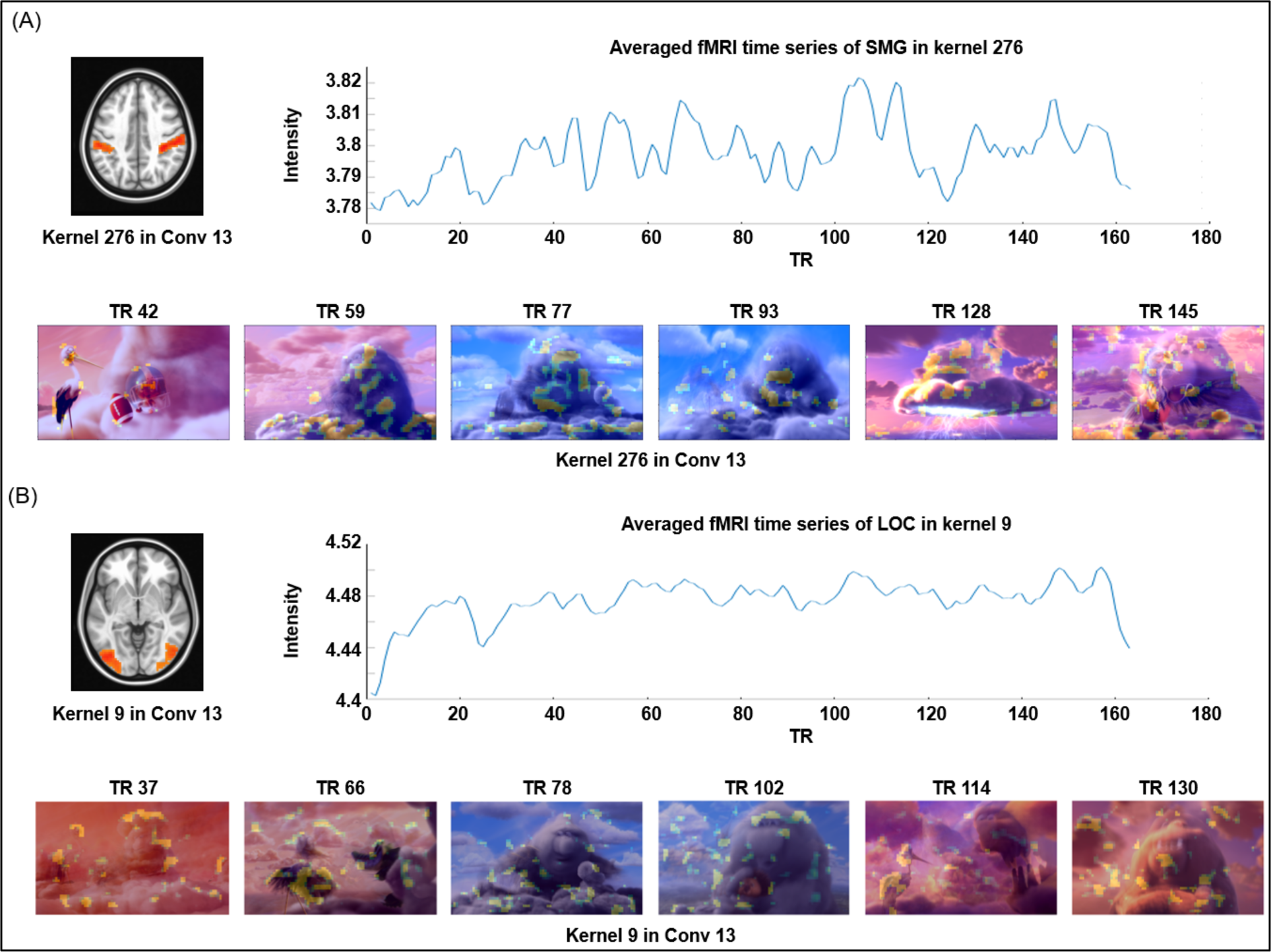
Reverse analysis of (A) supramarginal gyrus and (B) lateral occipital complex regions. Activation maps were obtained as GLM results for (A) kernel 276 for supramarginal gyrus and (B) kernel 9 for lateral occipital complex. The averaged fMRI time series are for each brain region, and the feature maps are for the TR times related to the peaks of the averaged fMRI time series.

The visual cortex emerged as the most predictable region associated with features from the 1^st^ convolutional layer. Specifically, we observed kernels 7, 43, and 63 of the 1^st^ convolutional layer showing activation only in the visual cortex (Figure 5A). Figure 5A showcases the BOLD time series in the visual cortex, allowing us to pinpoint the time points of peak regional activations. Furthermore, the feature activation maps corresponding to these peak time points for kernel 43 are also shown. These maps validate that the features associated with this particular kernel primarily emphasize edges or boundaries of objects, disregarding both the background and main characters.

The activations of the posterior cingulate cortex by features from the 1^st^ convolutional layer were somewhat unexpected. This region is involved in the default mode network, which is typically associated with higher-order brain functions. Nonetheless, we successfully identified kernels 6, 19, 24, 42, 55, and 58 of the 1^st^ convolutional layer that displayed activations only in the posterior cingulate cortex. Subsequently, we calculated the averaged BOLD time series within this region and identified the feature activation maps corresponding to time points exhibiting high BOLD activations (Figure 5B). Intriguingly, these maps indicate that the features within these kernels primarily relate to image backgrounds rather than the main characters.

The supramarginal gyrus displayed the highest probability of activations among all regions for the 13^th^ convolutional layer. Previous studies have associated this region with empathy for pain (Richardson et al., 2018). Within the supramarginal gyrus, we identified kernels 42, 103, 168, 276, 428, and 510 of the 13^th^ convolutional layer that exhibited activations. Subsequently, we extracted the time series of BOLD activations in this region (Figure 6A). The feature activations in these kernels often appear blurry due to earlier convolutions and max-pooling layers. To enhance interpretability, we overlaid the feature activations with the input image. Observing the overlaid images, it becomes evident that the feature activations may involve multiple characters (at time point 42) or be linked to facial expressions (at time point 145). These findings suggest that the features within these kernels possess the capacity to represent higher-order social information.

The lateral occipital complex region related to overall object shape perception was also noticeably observed in the 13^th^ convolutional layer. Unlike visual cortex, which is related to low-level visual elements, lateral occipital complex is associated with the task of comprehensively recognizing objects (Grill-Spector, 2001). Kernel 9, 43 and 128 in the 13^th^ convolutional layer were related to lateral occipital complex. After extracting the averaged time series of BOLD activations, feature maps from the peaks of the time series were analyzed (Figure 6B). Activation occurred across a variety of different objects (clouds, animals and background objects).

## 4. Discussion

In this study, we employed a widely used convolutional neural network, VGG-16, to extract diverse visual features in a movie, and associated these feature activations with brain activations recorded through fMRI. The observed brain activations demonstrated to some extent a hierarchical pattern across convolutional layers. Lower convolutional layers showed stronger associations with posterior visual cortex and posterior cingulate cortex, whereas higher convolutional layers exhibited stronger associations with lateral occipital cortex and supramarginal gyrus. However, within a given convolutional layer the differentiation of different feature activations was quite limited, leading to many different feature activations associated with similar brain regions.

As a sanity check, we first examined the brain activations associated with the first layer of VGG-16, where the features are thought to reflect most basic local features of an image (Krizhevsky et al., 2017; Zeiler & Fergus, 2014). As predicted, most of the associated brain activations were observed in the posterior occipital cortex, which corresponds to lower visual areas. The correspondence suggests, to some degree, similar visual features may be processed in shallow layers in the convolutional neural network and early visual brain areas. The current results also suggested limitations of using the temporal dynamic patterns to study brain activations. That is, the brain activation patterns from the different kernels of a layer were spatially highly similar and only showed a small number of distinct patterns. This makes sense given that the kernel activation time series from a layer were also highly correlated, and only a few principal components could explain most of the variance (Figure 2A). Even though different kernels extract distinct features, the changes of activations across different input may be highly similar, e.g., edges in different directions may produce highly correlated activations over time. The temporal smoothing with hemodynamic response function may also contribute to the high correlations. Nevertheless, the factors that cause temporal similarity in the convolutional neural network resemble those measured with BOLD fMRI, which also suffer from poor specificity in measuring visual responses in brain. Alternative approaches, such as multivoxel pattern analysis (Kriegeskorte et al., 2008) may be more effective to study the brain representation of different visual features.

Surprisingly, a small number of kernels in the first convolution layer were associated with brain activation in the posterior cingulate cortex, which is part of the default mode network (Raichle et al., 2001). The default mode network is thought to be situated in a higher hierarchy of organization (Margulies et al., 2016), and the associated functions are usually related to internalization and higher social functions (Buckner & DiNicola, 2019). However, recent studies also showed its involvements in naturalistic perception (Brandman et al., 2021). The current results provide further insights into task related activations in the default mode. Specifically, the current results demonstrated that the activations of visual features from the first layer of VGG-16 can be related to the activations in the posterior cingulate cortex. Feature analysis further indicated that the visual activations of these kernels were associated with the background, rather than the main characters of a scene. The activations in the posterior cingulate cortex may be related to the processing of unattended features in the background, or may be negatively correlated with the level of attentions (Kaefer et al., 2022). Nevertheless, the current results suggest that simple visual features may be linked to brain activity in the posterior cingulate cortex. Reverse inference of functions in the posterior cingulate cortex requires extra caution.

As the convolutional layer progresses deeper, the brain regions linked to kernel activations shift in a forward and upward direction within the brain. Specifically, there is a greater likelihood of association with the lateral occipital complex and the supramarginal network, both of which exhibit high inter-subject correlations (Di & Biswal, 2020). The lateral occipital complex plays a role in higher-order visual processing associated with object recognition (Grill-Spector et al., 2001), distinguishing itself from lower visual areas specialized in low-level visual features like texture (Malach et al., 1995). Hence, there appears to be a loose connection between the hierarchy of convolutional neural networks and the visual processing system in the brain. The reverse analysis further validated that feature activations linked to the lateral occipital complex were typically situated around the main characters or objects in a scene (Figure 6B). These findings align with a recent study demonstrating that responses from the lateral occipital complex were significantly predictive for scene, object, and action recognition in videos during movie viewing (McMahon et al., 2023).

The supramarginal gyrus serves as a crucial brain region for the recognition and comprehension of others’ emotions and pain (Di & Biswal, 2020; Lamm et al., 2011; Silani et al., 2013). The present findings indicate that details regarding a character’s intentions or pain experience may be encoded within a single frame of video stimuli, and these details can be extracted using convolutional neural network models. In the reverse analysis, the feature activations by the relevant 13th layer kernels may at times involve two characters and at other times be associated with a single main character (Figure 6A). Further investigations are needed to examine the activation characteristics of these 13th layer kernels across various images depicting social interactions, thereby validating our initial observations. Nonetheless, these results suggest that convolutional neural networks hold promise in representing advanced social information.

Several limitations in the current analysis should be taken into account. Firstly, the utilization of an animated video clip as a naturalistic stimulus might introduce a potential bias. Given its artificial nature, convolutional neural networks may be more predisposed to its specific characteristics. The transferability of such models to videos featuring real human actors remains an open and challenging question. Moreover, this study utilized a convolutional neural network to extract information from 2-D frames of a video, revealing that higher layers of VGG-16 may capture higher-order social information. However, social interactions often entail dynamic changes over time, and models considering temporal dependencies, such as recurrent neural networks, may be more apt for extracting such information. Nonetheless, the inclusion of temporal dependencies increases model complexity. Future investigations could explore alternative model options to enhance our comprehension of social interactions.

## 5. Conclusion

We have explored the utilization of a convolutional neural network model, specifically VGG-16, to extract diverse features at different levels. We linked these feature activations with brain activations measured through fMRI. Our analysis unveiled intricate relationships between brain activations and various layers of the convolutional neural network. Lower convolutional layers predominantly correlated with lower visual areas, while some exhibited connections with the posterior cingulate cortex, a component of the default mode network. In contrast, higher convolutional layers showed stronger associations with lateral occipital regions and the supramarginal gyrus, the latter being linked to higher-order social processing such as empathy for pain. Despite a high correlation in the temporal dynamics of kernel activations within the same layer, our findings suggest that kernels in deeper layers may signify more complex aspects of social interaction. The features extracted in this study provide a basis for future comparisons with alternative deep neural network models. Additionally, they hold promise for quantifying the content of movies and advancing our comprehension of the neural processes underlying cinematic experiences.

## Data and code availability statement

The fMRI data used in this study is public data sets and is available on openneuro (https://openneuro.org/; accession #: ds000228). The codes in this study are available upon a reasonable request to the corresponding author.

## Credit authorship contribution statement

**Wonbum Sohn**: Conceptualization, Formal analysis, Software, Investigation, Visualization, Writing–original draft. **Xin Di**: Conceptualization, Software, Funding acquisition, Project administration, Writing–review & editing. **Zhen Liang**: Writing-review & editing. **Zhiguo Zhang**: Writing-review & editing. **Bharat B. Biswal**: Supervision, Funding acquisition, Resources, Project administration.

## Data Availability

Data will be made available on request.

## Acknowledgements

This work was supported by (US) National Institute of Mental Health grants to Xin Di (R15MH125332) and Bharat B. Biswal (R01MH131335).

## References

Bartels, A., Zeki, S., & Logothetis, N. K. (2008). Natural Vision Reveals Regional Specialization to Local Motion and to Contrast-Invariant, Global Flow in the Human Brain. Cerebral Cortex, 18(3), 705– 717. 10.1093/cercor/bhm107

Brandman, T., Malach, R., & Simony, E. (2021). The surprising role of the default mode network in naturalistic perception. Communications Biology, 4(1), Article 1. 10.1038/s42003-020-01602-z

Buckner, R. L., & DiNicola, L. M. (2019). The brain’s default network: Updated anatomy, physiology and evolving insights. Nature Reviews Neuroscience, 20(10), Article 10. 10.1038/s41583-019-0212-7

Çelik, E., Keles, U., Kiremitçi, İ., Gallant, J. L., & Çukur, T. (2021). Cortical networks of dynamic scene category representation in the human brain. Cortex, 143, 127–147. 10.1016/j.cortex.2021.07.008

Chen, P.-H. A., Jolly, E., Cheong, J. H., & Chang, L. J. (2020). Intersubject representational similarity analysis reveals individual variations in affective experience when watching erotic movies. NeuroImage, 216, 116851. 10.1016/j.neuroimage.2020.116851

Deng, J., Dong, W., Socher, R., Li, L.-J., Li, K., & Fei-Fei, L. (2009). Imagenet: A large-scale hierarchical image database. 2009 IEEE Conference on Computer Vision and Pattern Recognition, 248–255. https://ieeexplore.ieee.org/abstract/document/5206848/

Di, X., & Biswal, B. B. (2015). Characterizations of resting-state modulatory interactions in the human brain. Journal of Neurophysiology, 114(5), 2785–2796. 10.1152/jn.00893.2014

Di, X., & Biswal, B. B. (2020). Intersubject consistent dynamic connectivity during natural vision revealed by functional MRI. NeuroImage, 216, 116698. 10.1016/j.neuroimage.2020.116698

Friston, K. J., Williams, S., Howard, R., Frackowiak, R. S. J., & Turner, R. (1996). Movement-Related effects in fMRI time-series: Movement Artifacts in fMRI. Magnetic Resonance in Medicine, 35(3), 346–355. 10.1002/mrm.1910350312

Grill-Spector, K., Kourtzi, Z., & Kanwisher, N. (2001). The lateral occipital complex and its role in object recognition. Vision Research, 41(10), 1409–1422. 10.1016/S0042-6989(01)00073-6

Hasson, U., Nir, Y., Levy, I., Fuhrmann, G., & Malach, R. (2004). Intersubject Synchronization of Cortical Activity During Natural Vision. Science, 303(5664), 1634–1640. 10.1126/science.1089506

Hu, W., Zhang, Z., Zhao, H., Zhang, L., Li, L., Huang, G., & Liang, Z. (2023). EEG microstate correlates of emotion dynamics and stimulation content during video watching. Cerebral Cortex, 33(3), 523–542.

Jiahui, G., Feilong, M., Visconti di Oleggio Castello, M., Nastase, S. A., Haxby, J. V., & Gobbini, M. I. (2022). Not so fast: Limited validity of deep convolutional neural networks as in silico models for human naturalistic face processing. Journal of Vision, 22(14), 3714. 10.1167/jov.22.14.3714

Kaefer, K., Stella, F., McNaughton, B. L., & Battaglia, F. P. (2022). Replay, the default mode network and the cascaded memory systems model. Nature Reviews Neuroscience, 23(10), Article 10. 10.1038/s41583-022-00620-6

Kriegeskorte, N., Mur, M., & Bandettini, P. (2008). Representational similarity analysis—Connecting the branches of systems neuroscience. Frontiers in Systems Neuroscience, 2. https://www.frontiersin.org/articles/10.3389/neuro.06.004.2008

Krizhevsky, A., Sutskever, I., & Hinton, G. E. (2017). ImageNet classification with deep convolutional neural networks. Communications of the ACM, 60(6), 84–90. 10.1145/3065386

Lamm, C., Decety, J., & Singer, T. (2011). Meta-analytic evidence for common and distinct neural networks associated with directly experienced pain and empathy for pain. Neuroimage, 54(3), 2492–2502.

Malach, R., Reppas, J. B., Benson, R. R., Kwong, K. K., Jiang, H., Kennedy, W. A., Ledden, P. J., Brady, T. J., Rosen, B. R., & Tootell, R. B. (1995). Object-related activity revealed by functional magnetic resonance imaging in human occipital cortex. Proceedings of the National Academy of Sciences, 92(18), 8135–8139. 10.1073/pnas.92.18.8135

Margulies, D. S., Ghosh, S. S., Goulas, A., Falkiewicz, M., Huntenburg, J. M., Langs, G., Bezgin, G., Eickhoff, S. B., Castellanos, F. X., Petrides, M., Jefferies, E., & Smallwood, J. (2016). Situating the default-mode network along a principal gradient of macroscale cortical organization. Proceedings of the National Academy of Sciences, 113(44), 12574–12579. 10.1073/pnas.1608282113

McMahon, E., Bonner, M. F., & Isik, L. (2023). Hierarchical organization of social action features along the lateral visual pathway. 10.31234/osf.io/x3avb

Nastase, S. A. (2019, May 16). Measuring shared responses across subjects using intersubject correlation | Social Cognitive and Affective Neuroscience | Oxford Academic. https://academic.oup.com/scan/article/14/6/667/5489905?login=true

Raichle, M. E., MacLeod, A. M., Snyder, A. Z., Powers, W. J., Gusnard, D. A., & Shulman, G. L. (2001). A default mode of brain function. Proceedings of the National Academy of Sciences of the United States of America, 98(2), 676–682. 10.1073/pnas.98.2.676

Rao, H., Wang, J., Tang, K., Pan, W., & Detre, J. A. (2007). Imaging brain activity during natural vision using CASL perfusion fMRI. Human Brain Mapping, 28(7), 593–601. 10.1002/hbm.20288

Raz, G., Winetraub, Y., Jacob, Y., Kinreich, S., Maron-Katz, A., Shaham, G., Podlipsky, I., Gilam, G., Soreq, E., & Hendler, T. (2012). Portraying emotions at their unfolding: A multilayered approach for probing dynamics of neural networks. NeuroImage, 60(2), 1448–1461. 10.1016/j.neuroimage.2011.12.084

Richardson, H., Lisandrelli, G., Riobueno-Naylor, A., & Saxe, R. (2018). Development of the social brain from age three to twelve years. Nature Communications, 9(1), Article 1. 10.1038/s41467-018-03399-2

Silani, G., Lamm, C., Ruff, C. C., & Singer, T. (2013). Right supramarginal gyrus is crucial to overcome emotional egocentricity bias in social judgments. Journal of Neuroscience, 33(39), 15466–15476.

Simonyan, K., & Zisserman, A. (2015). *Very Deep Convolutional Networks for Large-Scale Image Recognition* (arXiv:1409.1556). arXiv. http://arxiv.org/abs/1409.1556

Sun, Y., Ma, J., Huang, M., Yi, Y., Wang, Y., Gu, Y., Lin, Y., Li, L. M. W., & Dai, Z. (2022). Functional connectivity dynamics as a function of the fluctuation of tension during film watching. Brain Imaging and Behavior, 16(3), 1260–1274. 10.1007/s11682-021-00593-7

Zeiler, M. D., & Fergus, R. (2014). Visualizing and Understanding Convolutional Networks. In D. Fleet, T. Pajdla, B. Schiele, & T. Tuytelaars (Eds.), Computer Vision – ECCV 2014 (pp. 818–833). Springer International Publishing. 10.1007/978-3-319-10590-1_53

